# A framework for constructing insect steering circuits

**DOI:** 10.1101/2025.01.22.634241

**Authors:** Robert Mitchell, Barbara Webb

**Affiliations:** School of Informatics, The University of Edinburgh, Edinburgh, EH8 9AB, United Kingdom

## Abstract

Insects perform a variety of goal-directed navigation behaviours, each of which requires a comparison between their current and desired heading direction. Recent work has uncovered such a steering circuit in the fruit fly *Drosophila melanogaster*. We used the available neuroanatomical and physiological descriptions to derive six general rules which can be used to construct a class of steering circuits which operate in the same way. These rules are surprisingly permissive, suggesting that across insect species, steering circuits may have differing wiring while remaining functionally identical. We simulate an extreme example and demonstrate that it is functionally identical to the reported fruit fly circuit. Further, we argue that the principled approach we apply here could be applied more generally in performing comparative analyses across different insect species.

## 1 Introduction

Insects perform various goal-directed behaviours including maintaining a straight-line direction of motion [6], path integration and vector navigation [5], and long distance migration [15]. Each of these requires the animal to know the difference between the direction it is currently facing and the direction it wants to go. One way of obtaining this difference is to neurally encode both its current direction and the direction it wants to go (in a common spatial framework), then compare the two.

The insect brain region known as the central complex contains neural populations in which the location of a sinusoidal bump of activity tracks the current head direction of the animal, often described as an internal compass [16]. Here we take the term compass to mean that, over behaviourally relevant time scales and distances, the activity of each neuron can be consistently associated to the same azimuthal direction with respect to the surrounding environment (noting that this consistency need not necessarily persist across behavioural bouts). Recent work in the fruit fly *Drosophila melanogaster* [13] has uncovered a set of neurons in the central complex which appear to encode the animal’s goal direction with respect to its internal compass. As described in more detail in the following, these neurons modulate the relative activity of steering neurons which act in pairs to drive left/right turns that alter the heading direction of the animal, moving the phase of the compass bump to a specific (goal) position [13].

Here we analyse the anatomical model provided by Mussells Pires et al. [13] to derive a set of six rules, which can be used to construct a wide family of steering circuits that operate on the same principle through a combination of compass, goal, and steering neurons. Such rules are useful in producing bio-inspired models but also have relevance in comparative neuroanatomy; different insects may have steering circuits which are anatomically distinct but functionally identical. This is demonstrated by comparing selected steering circuits in simulation. We argue more broadly that such a principled approach to circuit analysis is useful in determining which features of the circuit are critical to its function.

## 2 Deriving the rules

### 2.1 Preliminaries

Here we will use vectors as an abstraction for neurons; discussion is limited to ℝ^2^ (2D with real components). In linear algebra, a basis is a set of vectors whose linear combinations can describe any point in space. For example, if

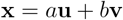

then we can choose *a* and *b* such that **x** can be any point, so long as **u** and **v** are linearly independent (i.e. not parallel).

A *positive basis* is an analogous concept in which the vector coefficients (*a* and *b* above) are restricted to positive values. This means that at least three vectors are required to span the plane (see figure 1). Now we have,

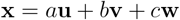

with *a, b, c* all strictly positive. In order to form a positive basis, the origin must lie within the convex hull of **u, v, w**. Equivalently, the inner angles between each adjacent vector pair must be *<* 180°, and the sum of the inner angles must equal 360°.

**Figure 1:**
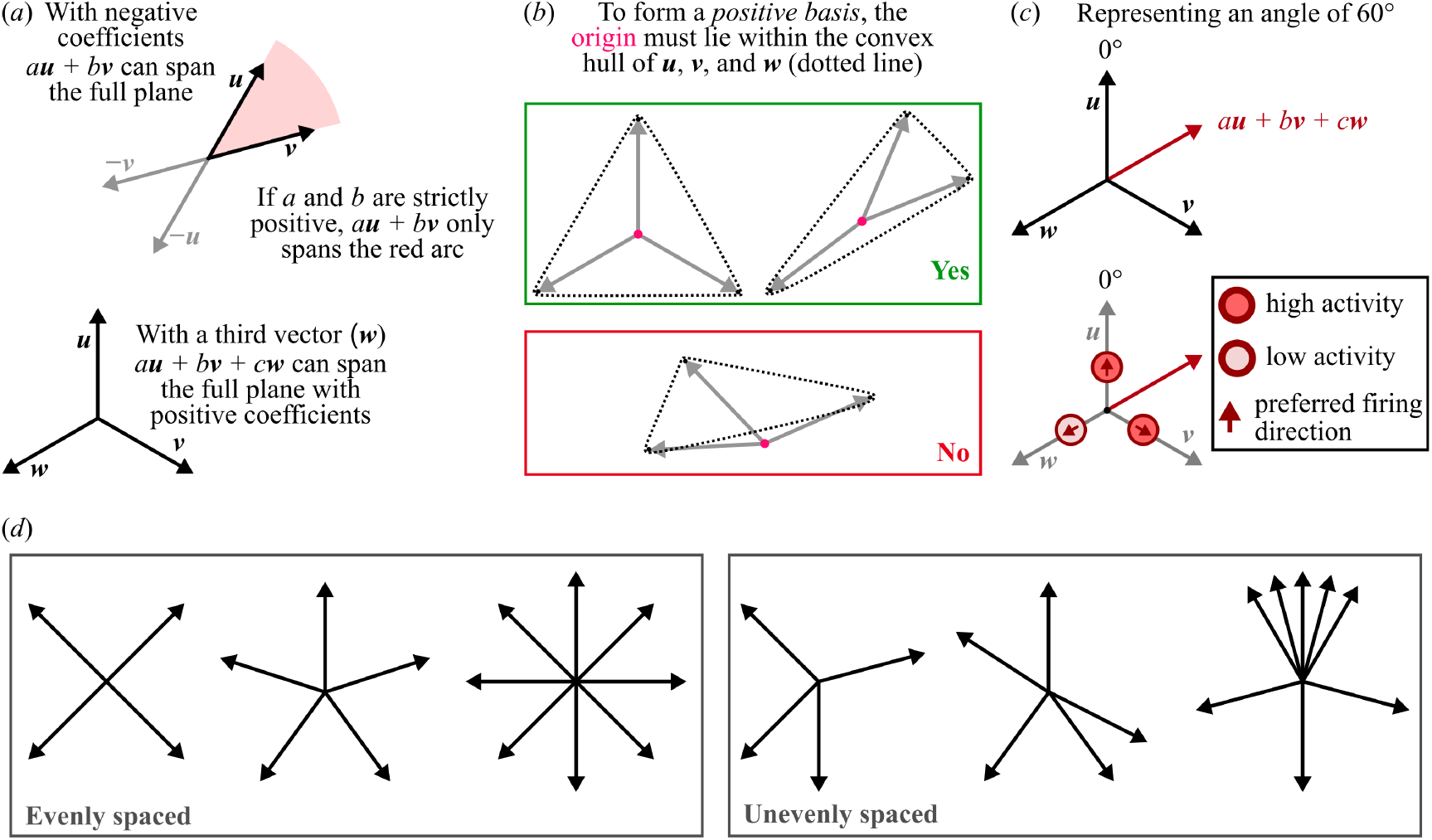
Positive bases. (*a*) Linear combinations of basis vectors can be used to span the plane. However, if negative values are prohibited (e.g. in a neural substrate), a third vector must be included to complete a *positive basis*. (*b*) To determine whether a set of vectors represents a positive basis, we can check whether the origin lies within the convex hull of the vectors. The examples in the green box both represent positive bases. The example in the red box cannot represent any angles which lie outwith the border vectors. (*c*) Translating the positive basis concept into a neural population. Basis vector coefficients are essentially translated into relative firing rates for the neurons which represent the basis vectors. So long as the vectors along the preferred directions of the population create a positive basis, the population can encode any angle. (*d*) Examples of valid neural bases with both even and uneven spacing. Note that the evenly spaced, four-vector example has no redundancy – removing any of the vectors will break the basis. In the uneven four-vector case, the two lower vectors are redundant, meaning either could be removed without breaking the basis. In the eight-vector example, uneven spacing could be exploited to increase the resolution in one part of the angular domain (for a neural representation).

The positive basis concept is illustrated in figure 1. The use of a positive basis to encode any angle is equivalent to the population vector average [3] commonly used to decode neural populations in the insect central complex (e.g. [16]). This assumes an angle is encoded as a vector which is decomposed across a population of neurons, each of which is sensitive to a different preferred direction. The neurons can be thought of as a set of vectors pointing along the preferred directions. The basic requirement for such a set to represent any angle is that it forms a positive basis.

Thinking of neural population codes in this way can be useful. For example, if we simply want to represent an angle, there is no requirement that neural tuning curves tile space evenly so long as a positive basis is formed. This means that, for example, a circuit could be robust to uneven spacing arising from errors in construction. Uneven spacing might also be useful: by improving redundancy in small circuits (figure 1d, compare evenly spaced and unevenly spaced four-vector examples); or increasing resolution in a certain part of the angular domain (figure 1d, unevenly spaced eight-vector example). Geometrically, the number and distribution of basis vectors used should make no difference to the precision, but biologically, it could [14].

### 2.2 A minimal circuit example

We will start by considering a minimal circuit and then generalise. The stages are illustrated in figure 2.

**Figure 2:**
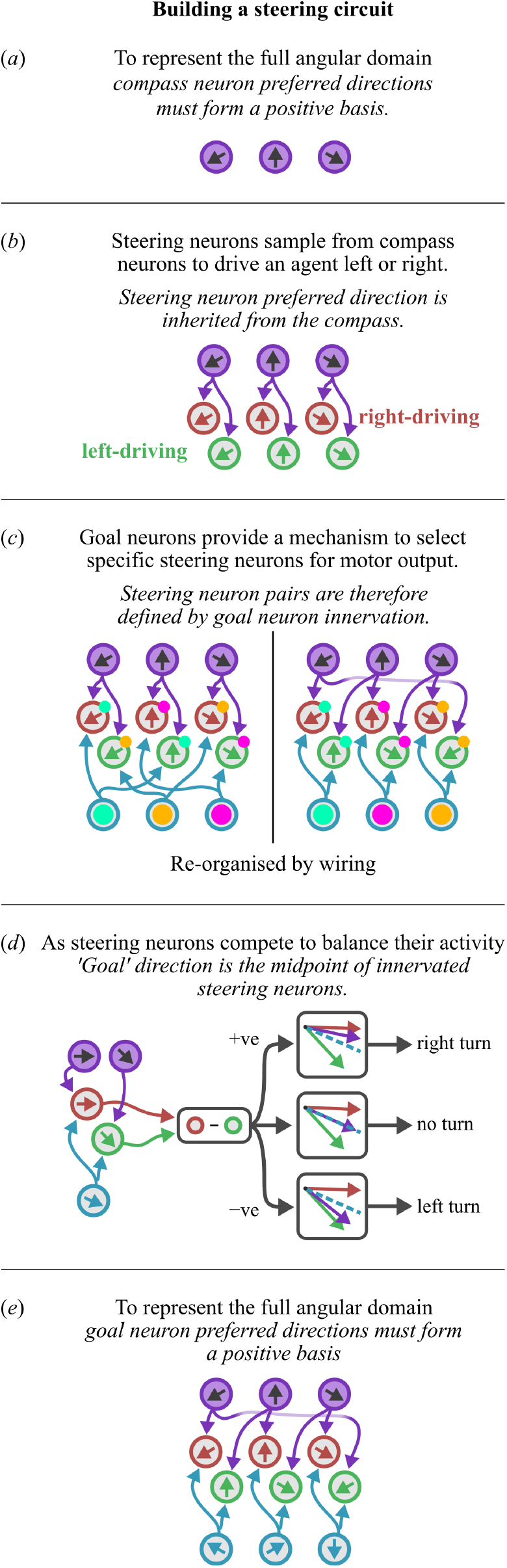
The circuit construction described in §2.2.

Compass neurons should be able to represent the full angular domain thus, *compass neuron preferred directions must form a positive basis*. The simplest example would be three neurons with evenly spaced receptive fields (figure 2a).

Following Mussells Pires et al. [13] (also see [10, 17]), steering neurons split into left-driving and right-driving populations, which steer the animal by rotating it to balance the activity between two selected points on the compass. In order to do this, steering neurons sample from the compass neurons meaning that *steering neurons inherit a preferred direction from the compass*. A simple steering population would have one left and one right steering neuron sampling from each compass neuron (figure 2b). We then require a way of selecting a left-driving and right-driving steering neuron to work as a pair to drive the agent towards their midpoint (note this implies that the steering neurons in our pair cannot connect to the same compass neuron).

The selection function is provided by goal neurons, each of which innervate at least one left-driving and one right-driving steering neuron. Thus *steering neurons are organised into pairs according to goal neuron innervation* (figure 2c). Further, this implies that *the ‘goal’ represented by a goal neuron is the midpoint of the innervated steering pair* (figure 2d), that is, the direction in which their activities will be balanced.

We now have a set of discrete goals which we can average to cover the continuous angular domain (as in the compass), therefore *the set of represented goals must form a positive basis*. We must choose goal neuron connections such that the set of possible goal directions creates a positive basis. In the simple example illustrated so far, this can be accomplished using three goal neurons, connected to steering pairs as shown in figure 2e.

One further observation arises from figure 2d. Steering neurons have both a driving direction (they drive left or right motor outputs) and a sampling direction (their preferred direction lies to the left or right of their combined goal). In figure 2d, right-driving neurons (red) are left-sampling and vice-versa. Whether right-driving neurons are right-sampling or left-sampling is not important as this simply defines whether they will steer away from the goal or towards it. But this means that *the relationship between steering and sampling should be consistent across all steering pairs*.

Our minimal example circuit is shown in figure 2e. It is neat and regular, however the observations we emphasise above generalise to allow circuits which are not neat and regular at all.

### 2.3 Generalised rules for steering circuits

In the following we will use vectors to represent directionally tuned neurons. We will refer to cells and the vectors representing those cells interchangeably; it can be easier to think in terms of a loose superposition of the two concepts (figure 1c).

A steering circuit is made up of three neural populations: compass, goal, and steering (which splits into *L* and *R*). For the purposes of construction, we can think of these populations as unit vector sets **C** for compass neurons, **G** for goal neurons, and **S**_*L*_, **S**_*R*_ for the two sets of steering neurons. Lower case sub-script refers to a specific neuron, e.g. **c**_*j*_ is the (vector describing the) *j*th compass neuron.

#### 1. C must form a positive basis

This means that any direction can be represented by some combination of compass neuron activity.

#### 2. Steering neuron direction is a weighted sum of the compass neurons sampled

In our simple example circuit, each steering neuron received input from one compass neuron, but in principle each could sample from multiple compass neurons in a weighted fashion. The direction of that steering neuron is then the weighted sum of all compass neurons sampled: **s**_*L,j*_ = *w*_1_**c**_1_ + *w*_2_**c**_2_ + *…* + *w*_*n*_**c**_*n*_ (assume **s**_*L,j*_ is then normalised, which also implies that ∥**s**_*L,j*_∥ ≠ 0). The sampled compass neurons do not need to be contiguous and different steering neurons could sample from different numbers of compass neurons. The sets of compass neurons sampled by different steering neurons may also overlap.

#### 3. Steering neurons form left/right pairs according to goal neuron innervation

Steering pair *j* is defined by the goal neuron **g**_*i*_ which provides input to those steering neurons. I.e. **s**_*L,j*_ and **s**_*R,j*_ both receive input from **g**_*j*_.
Note that steering pairs do not have to be exclusive. For example, one steering neuron could participate in multiple pairs so long as rules 4 and 5 are obeyed. This means that there could be different numbers of left and right steering neurons.

This also implies that goal neurons may innervate multiple steering neurons within the same *L* or *R* sub-population, forming two steering sets rather than strict pairs. The combined direction of one set could be considered the weighted sum of all neurons within that set (as in rule 2).

#### 4. The ‘goal’ represented by a goal neuron is defined by its innervated steering neurons

The steering mechanism works by selecting neurons from the compass layer and rotating the agent to balance their activity. Goal neuron input to steering neurons moderates their activity and acts as the selection mechanism. The represented goal can therefore be inferred by examining the connections from compass to steering neurons, and goal to steering neurons.

This inference process is described in detail in §5.3.2 and illustrated in figure 7. The process is general and includes the case where **s**_*L,j*_ and **s**_*R,j*_ actually represent weighted combinations instead of a pair (rule 3).

#### 5. G must form a positive basis

This provides that any desired goal direction can be represented by some combination of goal neuron activity.

#### 6. The sampling/driving relationship should be preserved across all steering pairs

Both **s**_*L,j*_ and **s**_*R,j*_ have a direction which is a weighted sum of their inputs. If **s**_*L,j*_ lies to the left of **s**_*R,j*_, then all *L* steering neuron directions must lie to the left of their paired *R* steering neuron directions. (It does not matter if left-driving steering neurons are left-sampling or right-sampling, so long as the relationship is the same across all pairs.) By the same logic, paired steering neurons cannot have the same direction.

Examples for each rule are given in figure 3, and an unintuitive example circuit is given in figure 4. We next consider whether such a strange example can be made to work.

**Figure 3:**
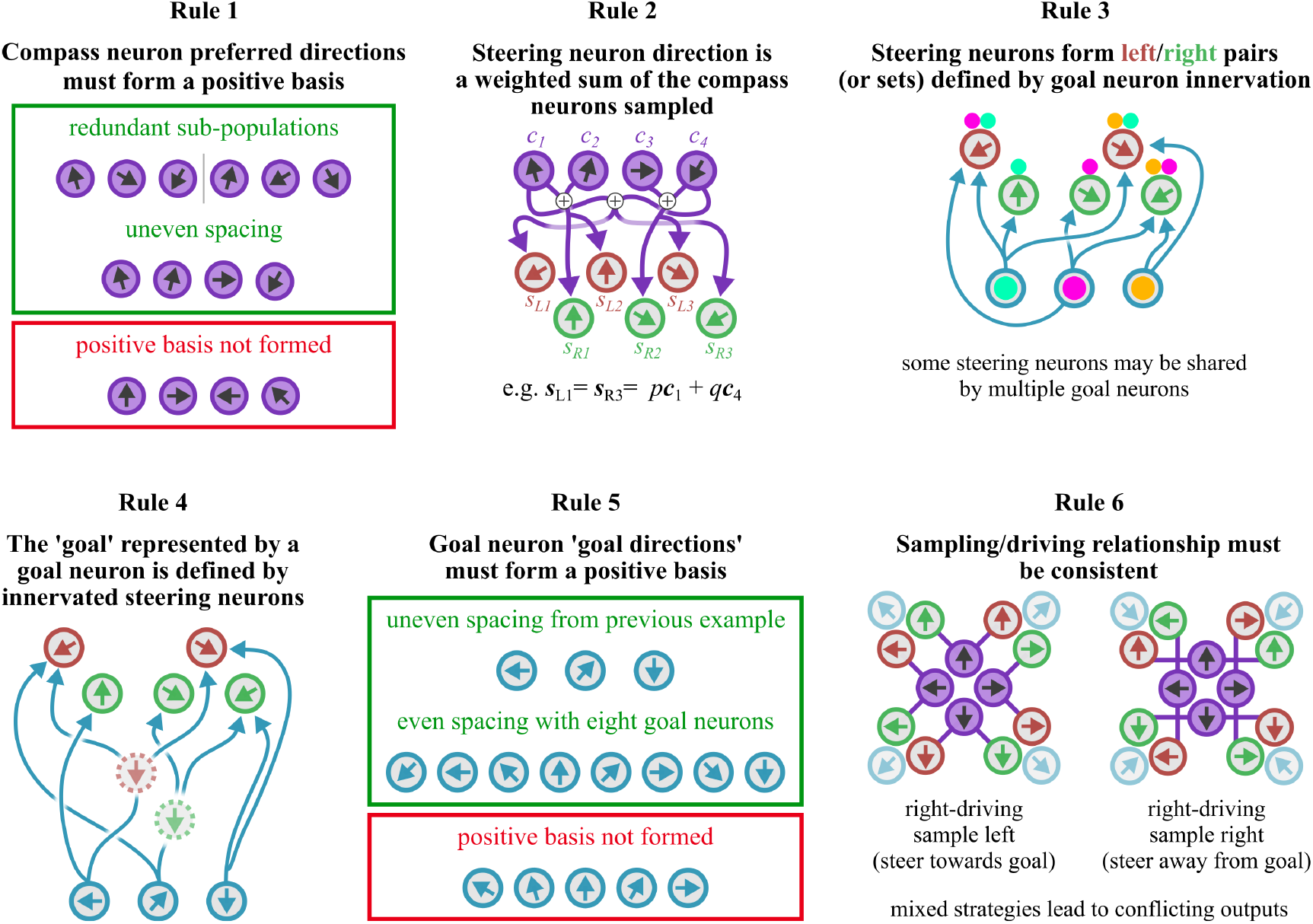
Illustrating rules for steering circuits. **Rule 1** Compass neuron preferred directions must form a positive basis (see figure 1). **Rule 2** Steering neuron direction is a weighted sum of the compass neurons sampled. In our example, we could choose weights *p* and *q* so that we can have any angle between the sampled compass neurons. **Rule 3** Steering neuron pairs or sets are defined by goal neuron innervation. Coloured markers indicate neurons which are grouped by goal neuron inputs. **Rule 4** The goal direction represented by a specific goal neuron is defined by the steering neurons it innervates. Where a goal neuron inputs to a set of steering neurons within the same steering group, these can be thought of as forming a virtual steering neuron whose direction is defined by a weighted combination of the set. This weighting could also be used to balance multiple neurons in one steering population against a single neuron in the other. **Rule 5** Resultant goal neuron goal directions must form a positive basis. **Rule 6** The sampling/driving relationship must be consistent. Either left-sampling neurons should drive right turns (towards goal), or left-sampling neurons should drive left-turns (away from goal). Mixed schemes would lead to conflicts between steering pairs.

**Figure 4:**
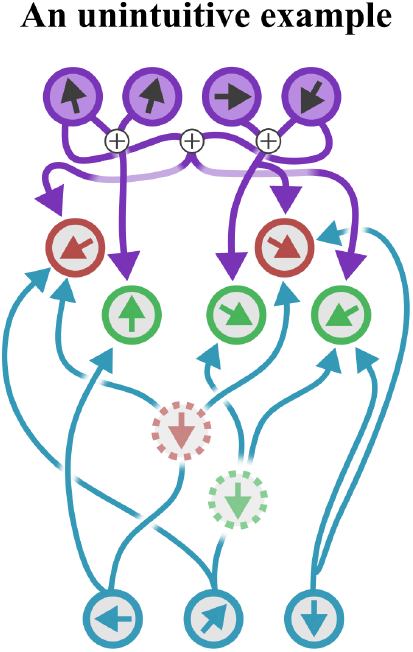
A hypothetical unintuitive example circuit. The compass population forms a positive basis with unevenly spaced tuning. Steering neurons receive input from compass neurons in a weighted fashion, such that two or three steering neurons are fed by four compass neurons. Left/right steering groups are of different sizes, meaning goal neurons must innervate sets of neurons in each side in order to generate a positive basis for goal neurons. The combined preferred direction for steering sets is indicated by a virtual steering neuron (dashed edge); this can be defined by weighting goal neuron connections onto steering neurons. This also implies that one could weight goal/steering neuron connections such that sets of neurons in one steering group can be balanced against a single neuron in the other. The angles between the steering pairs are also not consistent; goal neurons from left to right have steering pairs separated by 180°, 300°, and 120°. Note that 180° separation of a steering pair (virtual or otherwise) is permitted; this is an example where the vector sum analogy breaks down and we have to think about the mechanics of the circuit. Finally, the resultant goal directions are unevenly spaced but form a positive basis.

### 2.4 Demonstrating the functional equivalence of steering circuits

We created a simulation in which a model steering circuit is fed a series of goal angles, resulting in a random walk. We simulated the circuit from Mussells Pires et al. [13], our uniform circuit layout (with *n* = 3, 5, 8, and 21 neurons), and a version of our unintuitive circuit from figure 4.

We defined a circuit with the same neuron counts and connections as figure 4. We set the compass neuron preferred firing directions to 350°, 10°, 90°, and 200° (roughly as drawn). We then used stochastic optimisation (differential evolution) to find the weights for all present connections such that the steering output of the circuit matched (as closely as possible) that of the Mussells Pires et al. [13] circuit (figure 6). The resultant weights generated goal neuron preferred directions which did not match our hypothetical circuit in figure 4; in fact, the result was even more strange (figure 6c). Nevertheless, the circuit is clearly capable of performing the same function as the uniform circuits, and that of Mussells Pires et al. [13] (figure 5).

**Figure 5:**
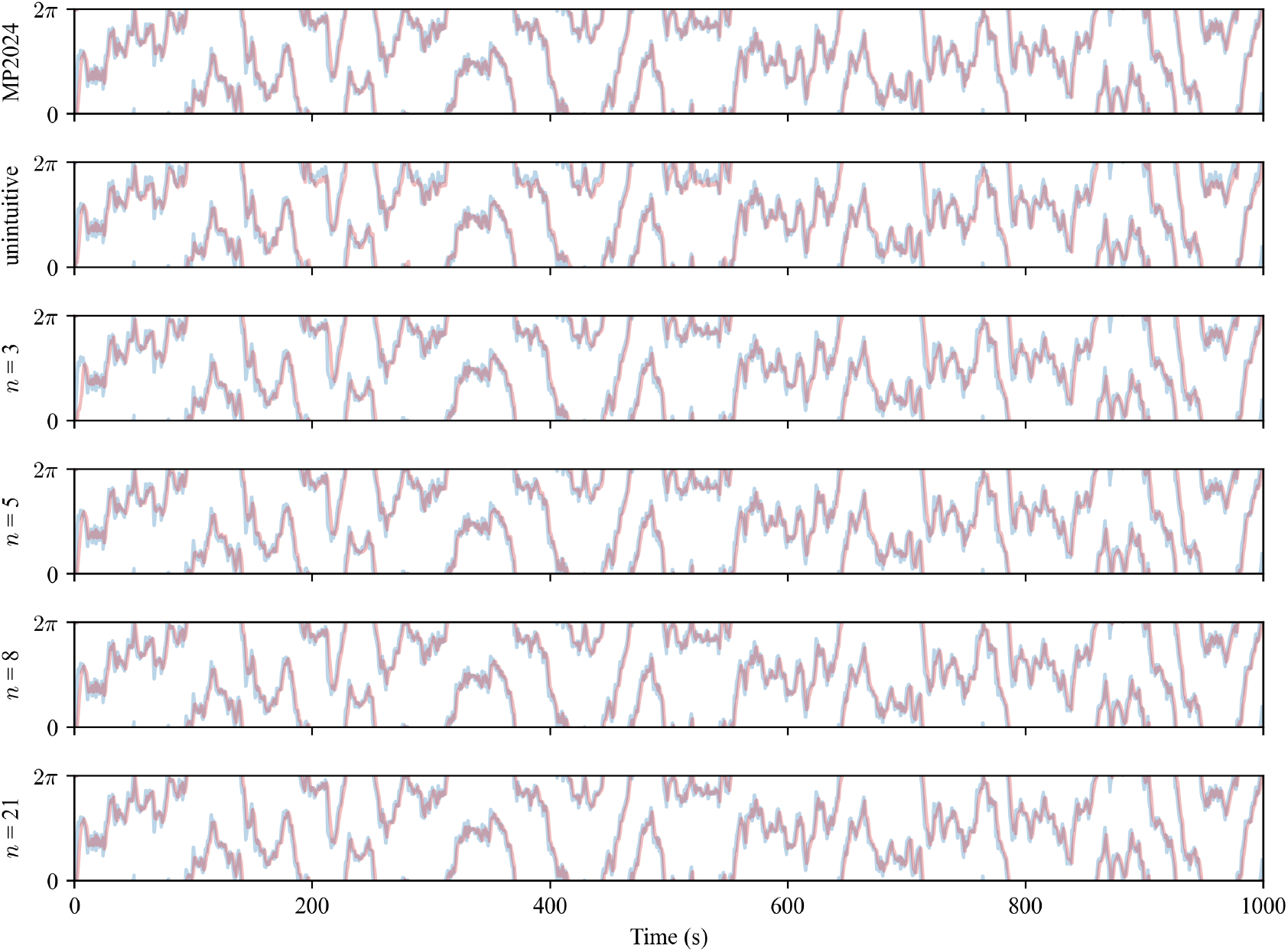
Random walk simulations. A random walk is generated by accumulating random samples from a von Mises distribution with *µ* = 0, 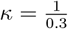. This random walk is then fed into the goal neurons for each circuit, and the circuit then steers to follow the walk. The blue line is the random walk over time, the red line is the direction steered by the agent over time. If the circuit is working properly, the lines should overlap. ‘MP2024’ is the circuit from Mussells Pires et al. [13], ‘unintuitive’ is the circuit from figure 6c (based on figure 4), and the remainder are circuits with uniformly spaced compass/goal neuron preferred directions with different numbers of neurons (*n* = 3 is the minimal circuit from figure 2e. All circuits are roughly tuned to steer at the same rate (between 50° and 60° per second). The *κ* parameter for the von Mises process was chosen such that the random walk would not change faster than the circuits could possibly follow. All circuits successfully follow the random walk.

**Figure 6:**
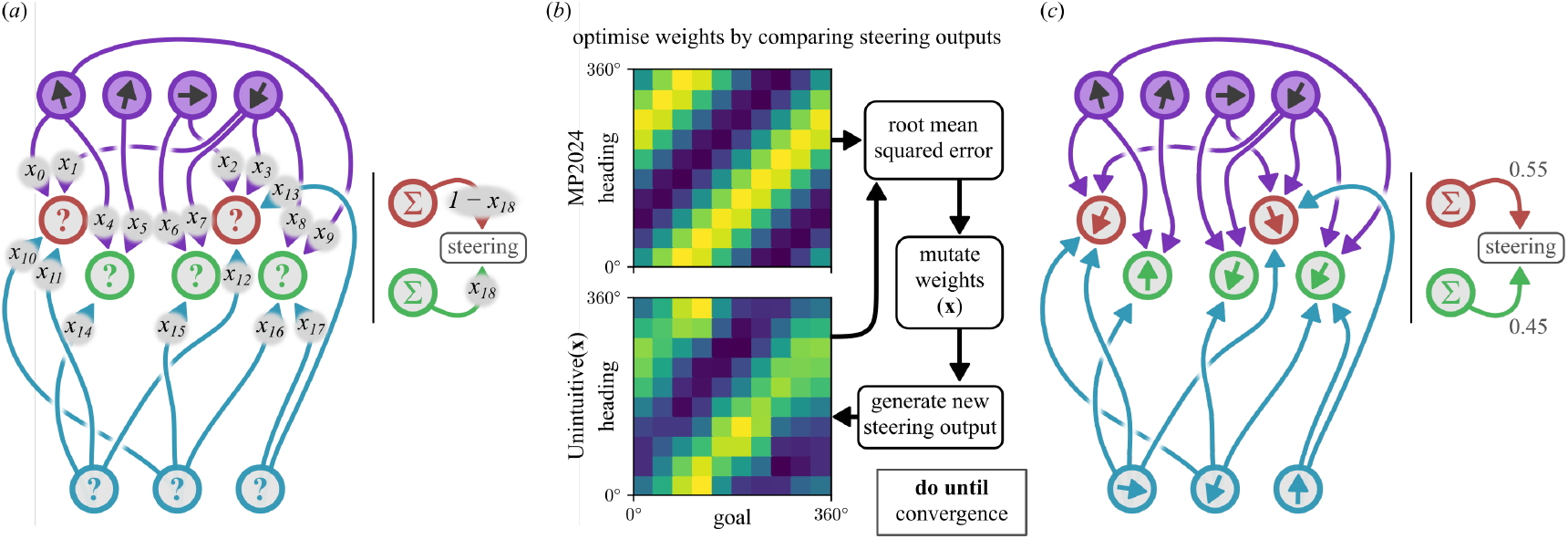
Creating a simulated version of the hypothetical circuit in figure 4 by comparing the steering output of a given parameterisation to the output from [13]. (*a*) The initial circuit is configured such that there are four compass neurons with preferred firing directions of 350°, 10°, 90°, and 200°. There are two *R* steering neurons (which steer left) and three *L* steering neurons (which steer right), and three goal direction neurons. A set of connections are chosen which may exist according to a parameter vector **x.** We also include a weight on the steering output meaning that (for example) *R* steering neurons may generate more steering than *L* neurons for the same level of activity. (*b*) The parameter selection process. We first generate the set of steering outputs from the Mussells Pires et al. [13] model (MP2024) by computing the steering signal for different heading/goal combinations (linearly sampled from 0° to 360°). A parameter vector is generated for the test circuit, then the steering output computed and compared to the model from [13]. The parameter vector is then mutated and the cycle continues until a minimum root mean squared error is achieved. The mutation process is driven by differential evolution, provided by [19]. (*c*) The final circuit generated by the optimsation process. Despite the strange steering neuron tunings, the goal direction neurons form a positive basis and the circuit can steer (figure 5).

## 3 Discussion

### 3.1 Rules for steering circuits

The rules laid out here allow a lot of flexibility in circuit construction. Some examples:

- The limitations on how steering neurons sample compass neurons imposed by these rules are consistent with a very broad variety of sampling patterns, including uneven and overlapping sampling. The resultant angle between steering neuron directions could even be different for different steering pairs, so long as rules 6 and 5 are obeyed.
- Rule 5 implies that ‘represented goals’ do not have to be equally spaced, and similarly, rule 1 implies that compass neuron tuning does not have to be equally spaced.
- There is no requirement placed on the number of neurons in each layer. Steering neurons can be shared across different steering pairs so long as rules 6 and 5 are obeyed.

As described in figure 2, a minimal but sufficient circuit could be constructed with three compass neurons, three goal neurons, and six steering neurons (figure 2e). Interestingly, the minimal circuit is very similar to a model circuit evolved by Haferlach et al. [10]. The only real difference is that their circuit includes speed inputs to both integrate and discharge a homing vector for path integration. In terms of steering principle, they are identical.

The minimal circuit implies that the steering circuit anatomy described for the fruit fly [13] is more complicated than it needs to be. This immediately leads us to wonder why. While all circuits consistent with the stated rules may operate in the same way, they may produce subtle (or not so subtle) variations in behaviour or dynamics.

For example, recent work has shown there is a trade-off between the number of computational units in a (ring attractor) compass circuit, and the precision of tuning required for accurate angular representation [14]. Anecdotally, we observed that uniform steering circuits with larger numbers of neurons were more robust to changes in the neural activation function. Thus, while it is possible in principle to have a compass made up of three neurons, this could be more error-prone or more difficult to tune than than one with say, sixteen neurons. It is also worth noting that a three neuron network (or more generally, circuitry spanning the midline with an odd number of neural computational units) may be less likely to evolve (or harder to produce during development) in bi-lateral animals. Constraints on ancillary connections, such as the ring attractor architecture of the compass in insects, may also place restrictions on the number of compass neurons under certain assumptions [1].

Thus, circuit variation could be behaviourally relevant (e.g. improving some aspect relevant to the ecology of a specific animal). Or it could be the result of evolutionary and/or developmental constraints with minimal behavioural impact. But critically, large differences in morphology (wiring) or physiology (e.g. spread in preferred tunings) may not change how the circuit actually works (compare our unintuitive model in figure 6c with the anatomical model from [13]).

More generally, this principled (or rule-based) approach to analysing neuromorphology and physiology could be useful in filtering out which properties are functionally relevant. Two simple examples can be found in the fruit fly circuit described by Mussells Pires et al. [13]:

- There are cases where one steering neuron samples from multiple compass neurons or where different steering neurons sample from the same compass neurons. This multiple sampling is distinctive and appears to be functionally important. Applying our rules, the specific sampling pattern is not functionally relevant so long as rules 2, 6, and 5 are followed.
- The compass neurons in their model split into two overlapping neural populations (following the anatomy of the fruit fly). Most preferred directions are represented twice over in the population and some steering neurons sample from different sub-populations. Again, this is a fairly distinct anatomical feature and it is not clear if it is functionally relevant. Rules 3, 2, and 6 tell us how steering neuron pairs are defined, and how steering neurons must sample from compass neurons. Rules 4 and 5 tell us how we must wire up the goal neurons. Finally, rule 1 tells us that the double representation of preferred directions does not affect the function of the steering circuit.

It is useful to know where there are differences in anatomy and we should highlight that the relevance of a given anatomical feature depends on the function under scrutiny. While the double-representation of compass neurons is not important for the steering circuit it could be important elsewhere. For example, in the fruit fly it appears to subserve the mechanism by which self-rotation inputs shift the compass bump. That said, it is equally important to consider the underlying operational principles. Where our models work, we know we have a grasp on the underlying algorithm; where they do not, we have found a relevant anatomical difference.

### 3.2 Goal neurons and changing reference frames

Tangentially, it is worth considering how we define a ‘preferred direction’ for goal neurons. When modelling neurons which represent angles as a sinusoidal population code, the typical construction for a given neuron is:

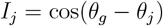

where *I*_*j*_ is the input to the *j*th goal neuron, *θ*_*g*_ is the goal direction and *θ*_*j*_ is the preferred direction of this specific neuron (there may be other set dressing around this cosine, but this is the core). To define the model population we must then define preferred directions for each neuron in the population, but deceptively, these preferred directions are not anchored to the external world.

While we may model a neuron as having a preferred direction of 0°, what this actually represents in terms of behaviour depends on the steering neurons this goal neuron innervates, which subsequently depends on the compass neurons sampled by those steering neurons. Thus, even in a circuit with a perfect geocentric compass, the preferred directions we give the goal neurons are arbitrary, unless we analyse the wiring to choose anatomically defined preferred directions [13]. Further, in insects, the inputs to the compass are plastic [8, 11, 9] which (in theory) allows the compass representation to drift or otherwise change with respect to the world. The same pattern of goal neuron activity could represent two different geocentric directions, depending on how well calibrated the compass is.

This may not matter for brief goal-directed behaviours such as straight-line orientation [6] (in fact, it could be beneficial if your aim is to be unpredictable [4, 2]). However, for an animal which performs vector navigation to a set location [17, 12], it is critical for the compass to stay consistent with respect to the world for the duration of the behaviour. How the insect stays consistent is, at present, unknown. The compass does have a calibration mechanism [7], however this mechanism does not seem sufficient to eliminate all compass drift.

## 4 Conclusions

Following the work of Mussells Pires et al. [13], we have presented here a set of general rules according to which insect-inspired steering circuits may be constructed. While these rules were derived for modelling purposes, we believe that they may be useful in analysing and comparing real steering circuits. More generally, we believe that this principled (or rule-based) approach to analysing insect neuroanatomy is valuable in distinguishing which morphological characteristics of a circuit make it functionally distinct.

## 5 Methods

### 5.1 Circuit simulations

#### 5.1.1 Rate model

Neurons are represented using a simple firing rate model.

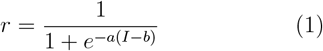

where *r* is the firing rate of the neuron and *I* is the input. The slope and bias parameters are set to *a* = 2 and *b* = 0.6 respectively. The parameters were tuned by hand and the same activation function was used for all neuron types and across different models. A bold **r** is used to denote a vector of firing rates (e.g. **r**_*C*_ is population response for the compass neurons).

#### 5.1.2 Compass neurons

Compass neuron input is

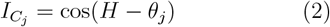

where *H* is the agent’s current heading and *θ*_*j*_ is the preferred firing direction (pfd) for the *j*th neuron.

#### 5.1.3 Goal direction neurons

Goal direction neuron input is given by

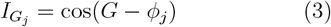

where *G* is the agent’s current goal direction and *ϕ*_*j*_ is the preferred direction of the *j*th goal neuron.

#### 5.1.4 Steering neurons and steering commands

Steering neuron input is

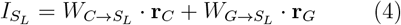

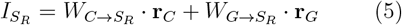

where *W*_*P →Q*_ represents the weight matrix from neural population *P* to population *Q*. The final steering command is given by

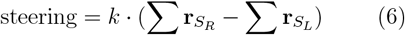

where *k* is a scaling parameter which determines how quickly the network can steer.

### 5.2 Uniform circuits

Uniform circuits are made up of *N* compass neurons, *N* goal direction neurons, and 2*N* steering neurons. Compass neuron preferred directions are evenly spaced over 360°

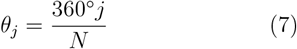

where *j* ranges from 0 to *N* − 1. Steering neurons receive input from compass neurons as

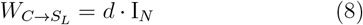

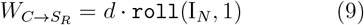

where I_*N*_ is the identity matrix of size *N*, `roll`(*X, n*) is NumPy’s roll operation (which shifts and wraps the columns matrix *X* by *n* positions), and *d* = 0.2 is a scale parameter which determines how much influence the compass neurons have over the steering neurons.

Goal neurons provide input to steering neurons according to

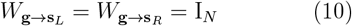

Steering neurons are thus organised into pairs which receive input from neighbouring compass neurons.

#### 5.2.1 Goal neuron preferred firing direction

The evenly spaced compass neuron tunings and structured columnar input to the steering neurons make it relatively straightforward to infer the goal neuron tuning. The angular tuning for compass neuron *j* can be expressed as a complex number

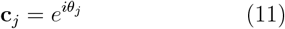

The tunings for each of the steering neurons are then simply

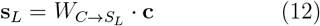

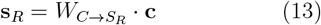

For these circuits, goal neuron preferred directions lie at the midpoint of the preferred directions of the innervated steering pair, so we first work out the corresponding left and right components of the goal neuron preferred direction, then we can sum these to get the final vector of goal neuron preferred directions.

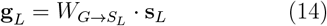

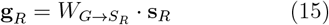

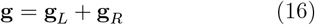

Then, remembering that the elements of **g** are complex numbers, *ϕ*_*j*_ in equation 3 is given by the complex argument of **g**_*j*_.

### 5.3 Unintuitive circuit

The unintuitive circuit contained four compass neurons, two L steering neurons, three R steering neurons, and three goal neurons. Compass neuron preferred directions were chosen arbitrarily as {350°, 10°, 90°, 200°} (unevenly spaced but obeying rule 1). Weight matrices were chosen such that some connections definitely did not exist and others could exist with a weight to be determined by an optimisation procedure.

Compass neurons input to steering neurons according to

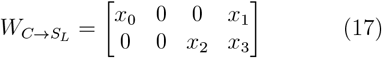

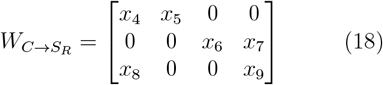

Goal neurons input to steering neurons according to

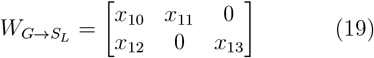

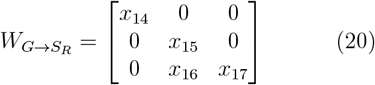

Thus, steering neuron preferred directions and by extension goal neuron preferred directions are unknown until these weights are provided. Finally, we modified the steering output (equation 6) to include a weight parameter

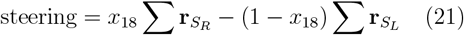

Functionally, this allows the point at which steering neuron activity is balanced to be tuned such that it does not lie exactly at the midpoint (see figure 7). The inclusion of this parameter is discussed in §5.3.2.

**Figure 7:**
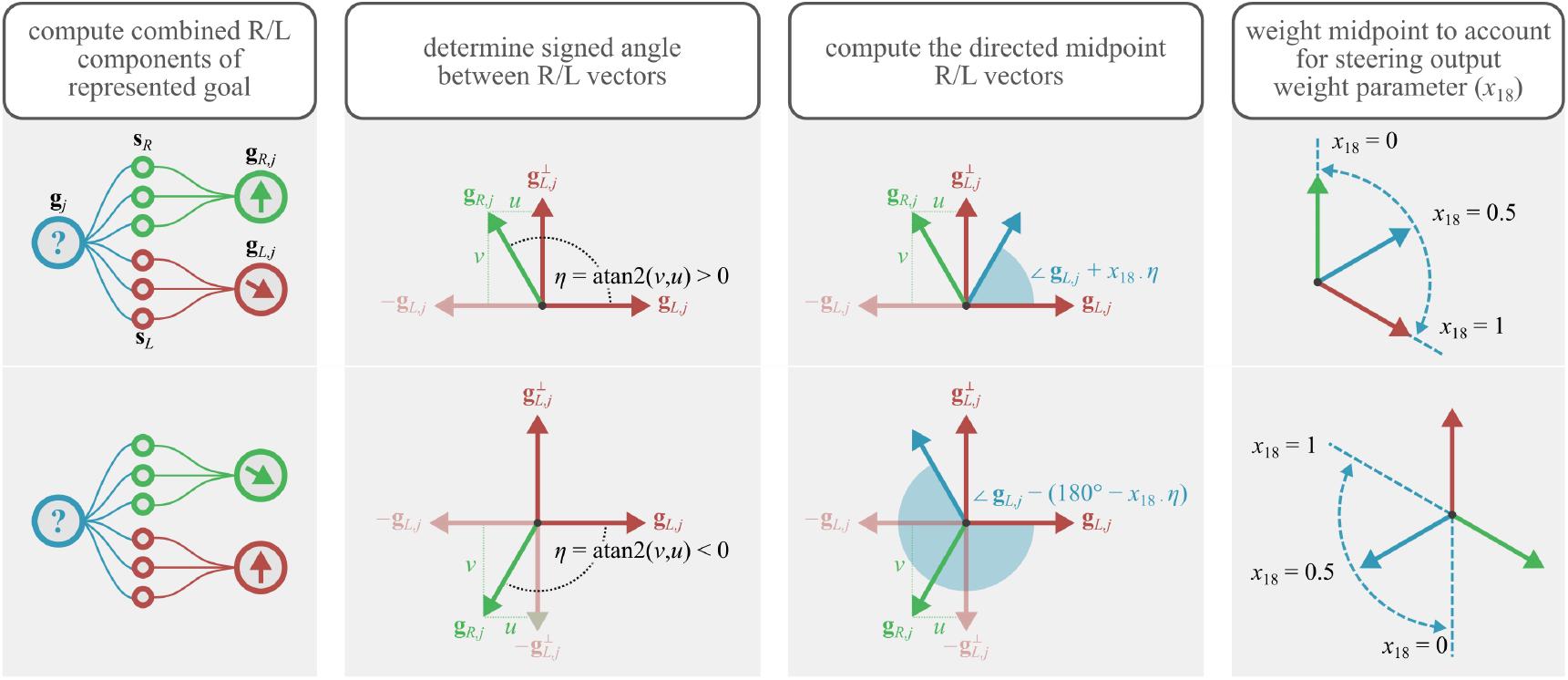
Visualisation of the goal direction neuron tuning inference. The first stage is to compute the combined steering direction for all steering neurons, on the same side, innervated by a goal neuron (equations 14 and 15). We then compute the signed angle (*η*) between the right component and the left component. The directed midpoint is then computed differently depending on whether the signed angle is positive or negative. A positive result indicates that we should use the midpoint of the inner angle, a negative result should use the outer angle, to respect the directionality of the steering neurons. Finally, this midpoint is weighted by model parameter *x*_18_ (equation 21), to reflect the bias introduced to the goal neurons by the weighted steering output. Rule 1 is guaranteed as we chose the compass neuron tuning, rules 2, 3, 4, and 6 define the goal tuning inference process, and rule 5 is obeyed by the resultant circuit.

#### 5.3.1 Optimisation procedure

In order to find a suitable parameter vector **x**, we linearly sampled angles from 0° to 360° and computed the steering output for every heading/goal combination, using the model from Mussells Pires et al. [13]. For a given parameterisation of the unintuitive circuit, we would compute the set of all possible steering outputs in the same way and compute the root mean squared error against the output from Mussells Pires et al. [13]. This root mean squared error was minimised using differential evolution [19, 18]. Parameters *x*_0_-*x*_17_ were permitted to vary between 0 and 2, and *x*_18_ was permitted to vary between 0 and 1. The bounds for *x*_0_-*x*_17_ were chosen arbitrarily (though positive as these connections appear to be excitatory) and those for *x*_18_ were dictated by equation 21. The initial state of the network, the optimisation process, and the result are shown in figure 6.

#### 5.3.2 Goal direction neuron inference

Inferring the goal neuron direction is complicated by the variability in steering neuron sampling of compass neurons, and goal neuron innervation of steering neurons.

Steering neuron preferred directions are computed as in equation 12 and 13; however in this case, the resultant vectors are all normalised. Similarly, the left and right components of the goal neuron directions are computed using equations 14 and 15, with the results being normalised. The normalisation is required as the lengths of the preferred direction vectors can vary, as they are made up of an arbitrary weighted vector sum; subsequent computations assume unit vectors (e.g. figure 7).

As we cannot make any assumptions about the angle between the left and right components of the goal, the simple vector sum used previously is not sufficient. In the case where the angle between the left and right components is greater than 180°, the represented goal will point in the wrong direction, and if the angle is equal to 180° then the vector sum will give a zero vector. We therefore have to account for the sign of the angle between the left and right components, and work with the angles directly instead of using vectors. A visualisation is given by figure 7.

For a given goal neuron *j*, combined left-hand component is given by **g**_*L,j*_ and the right-hand component by **g**_*R,j*_ (equations 14 and 15). We begin by computing 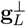, the set of left-hand component vectors which are orthogonal (counter-clockwise) to their counterparts in **g**_*L*_. We then compute the projection of **g**_*R,j*_ onto **g**_*L,j*_ and 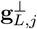.

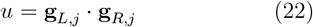

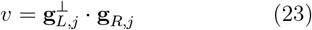

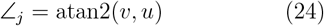

The final stage gives the signed angle ∠_*j*_ between **g**_*L*_ and **g**_*R*_. In our implementation, *L* steering neurons drive the agent right, and *R* steering neurons drive the agent left. Thus, the final goal preferred direction can be computed as

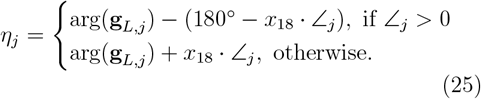

and to remain consistent with our complex/vector representation

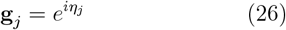

If we limit *x*_18_ to be 0.5, the process amounts to choosing **g**_*j*_ or *−***g**_*j*_, depending on the relative position of **g**_*L,j*_ and **g**_*R,j*_. If we let *x*_18_ vary, this allows the optimisation procedure to choose where the goal lies between **g**_*L,j*_ and **g**_*R,j*_ (figure 7).

As the optimisation procedure is stochastic, it is not possible to concretely state whether or not this extra parameter is necessary. However, it did produce better results with less optimisation attempts. We would therefore tentatively say that, where there are different numbers of steering neurons in each *L*/*R* sub-population, it is likely that this needs to be counterbalanced by weighting motor output. This may be an additional developmental constraint in real insect brains which enforces a degree of symmetry.

### 5.4 Steering model from Mussells Pires et al. [13]

The Mussells Pires et al. [13] model is implemented using the description given in their paper. Goal (FC2) neuron preferred directions are: −15°, −35°, −75°, −105°, −135°, −165°, 165°, 135°, 105°, 75°, 45°, 15°. *L* steering (PFL3_*L*_) neuron preferred directions are: 67.5°, 22.5°, *−*22.5°, *−*22.5°, *−*67.5°, *−*112.5°, *−*112.5°, *−*157.5°, 157.5°, 157.5°, 112.5°, 67.5°. *R* steering (PFL3_*R*_) neuron preferred directions are: *−*67.5°, *−*112.5°, *−*157.5°, *−*157.5°, 157.5°, 112.5°, 112.5°, 67.5°, 22.5°, 22.5°, *−*22.5°, *−*67.5°. The response for the *j*th steering neuron (in either the *L* or *R* population) is given by

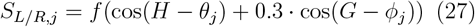

where *H* is the current heading *θ*_*j*_ is the preferred direction for the *j*th steering neuron, *G* is the goal direction, and *ϕ*_*j*_ the preferred direction for the *j*th goal neuron. The function *f* is

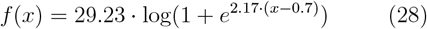

as in [13]. The final steering response is

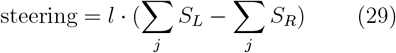

The constant *l* = 0.00018 is used to scale the model output so that it operates on roughly the same timescale as our other models.

## 6 Code availability

Our code is available on Zenodo (https://doi.org/10.5281/zenodo.14713161).

## 7 Author contributions

RM: conceptualization, investigation, methodology, software, validation, visualization, writing - original draft preparation, writing - review and editing. BW: conceptualization, funding acquisition, methodology, project administration, supervision, writing - review and editing.

## 9 Acknowledgements

We would like to thank the members of the InsectRobotics lab who provided feedback on early versions of this work. In particular, we would like to acknowledge the contributions of Yihe Lu and Evripidis Gkanias. We would also like to thank Auguste de Pennart and Stanley Heinze of the Lund Vision Group for their valuable feedback on the work.

